# Hepatocyte Embryonic Ectoderm Development (Eed) Deficiency Causes Liver Injury, Fibrosis, and Impacts Liver Regeneration

**DOI:** 10.64898/2026.03.13.711572

**Authors:** Yousra Ajouaou, James Griffin, Sade Chaffatt, Charlene Chen, Balnur Ibrash, Martine McManus, Kirsten C. Sadler

**Affiliations:** Program in Biology, NYU Abu Dhabi, PO Box 129188, Abu Dhabi, United Arab Emirates; Center for Genomics and Systems Biology, NYU Abu Dhabi, PO Box 129188, Abu Dhabi, United Arab Emirates

**Keywords:** Liver regeneration, EED, H3K27me3, epigenetics, PRC2, mice, fibrosis

## Abstract

Regeneration depends on tightly coordinated transcriptional programs governed by a dynamic epigenetic landscape to regulate cell identity, proliferation, and tissue remodelling following injury. The liver is highly regenerative due to its ability to rapidly upregulate genes that drive the cell cycle and other genes important for hepatocyte proliferation. In the uninjured liver, many of these genes are poised for activation based on promoter occupancy of trimethylated histone 3 lysine 27 (H3K27me3) and regenerative stimulus provided by partial hepatectomy leads to evacuation of this mark, suggesting this is a key component of proregenerative gene regulation in the liver. H3K27me3 is deposited by the polycomb repressive complex 2 (PRC2). Were we deplete H3K27me3 using hepatocyte specific deletion of key component of PRC2, Embryonic Ectoderm Development *(Eed^HepKO^*). This results in reduced liver size, increased hepatocyte death, proliferation, and fibrosis associated with upregulation of cell cycle and fibrogenic genes. Though these mice are less likely to survive two-thirds partial hepatectomy than WT (WT) controls, those that do survive increase liver mass faster than WTs. Importantly, genes occupied by H3K27me3 in uninjured WT livers are upregulated in EED^HepKO^ livers, and become further induced following PH. This demonstrates that PRC2 loss induces liver injury and dysregulates pro-regenerative gene expression, highlighting that epigenetic modulation depends requires precise targeting to achieve enhanced regenerative

## Introduction

Regenerating lost or damaged tissue is essential for all organisms to successfully mitigate injury. The field of regenerative medicine aims to promote regeneration by understanding and manipulating the cellular and molecular processes that govern tissue repair and regrowth. Epigenetic regulation of pro-regenerative genes is an attractive target to achieve these goals since epigenetic marks are reversible and can be altered by manipulating the complexes that write and erase them (1, 2). To achieve this, it is important to determine which epigenetic modifications are the most important for regulating regeneration relevant genes.

While most tissues in mammals cannot regenerate, the liver is a notable exception. Following two-thirds partial hepatectomy (PH), or modest hepatocyte injury, the remaining hepatocytes reenter the cell cycle to restore liver mass (3). This process is dictated by a highly coordinated pattern of transcriptional changes that culminate in the synchronous progression of hepatocytes through the cell cycle. A leading hypothesis is that the uninjured liver, as well as other regeneration competent tissues, have an epigenetic prepattern which allows the pro-regenerative genes to be activated quickly in a precise, coordinated pattern following loss of liver mass or function. This is supported by recent studies from our group (4) and others (5–7) showing that in uninjured, regeneration-competent tissues, pro-regenerative genes are in open chromatin bivalently marked by the repressive modification histone H3 lysine 27 trimethylated (H3K27me3) and activating mark, H3K4me3. We found that H3K27me3 is lost from some of these genes during regeneration (4) and others suggest this mark is important for regeneration competency in the liver (8). Additionally, young mice with deletion of the epigenetic regulator ubiquitin, PHD, and RING finger domain containing 1 (*Uhrf1*) in hepatocytes (*Uhrf1^HepKO^*) have accelerated activation of proregenerative genes following PH and accelerated liver regeneration associated with global loss of H3K27me3 from promoters of proregenerative genes (9). Together, these support the hypothesis that in hepatocytes, PRC2-mediated H3K27me3 occupancy of cell cycle and other pro-regenerative genes serves as the gatekeeper to their activation following liver injury

Previous studies have investigated the role of H3K27me3 in the liver via deletion of the PRC2 enzymatic factors, enhancer of zeste 1 or 2 (Ezh1, Ezh2), alone or in combination in hepatocytes showing that there is redundancy in their function (10–13). The exception is when Ezh2 is knocked out in hepatoblasts, which effectively depletes H3K27me3 and promotes acquisition of biliary identity (14) and reduces liver size (10). While Ezh1 knockout mice have no discernible phenotype (15) and Ezh2 knockout is embryonic lethal (16), to study the function of PRC2 in the adult liver, investigators have used Ezh1 full-body knockouts (*Ezh1 ^−/−^*) with hepatocyte-specific knockout of Ezh2 Cre under the albumin promoter (Alb:Cre; *Ezh1 ^−/−^/Ezh2^HepKO^*). These mice are viable; however, they develop multiple liver defects, including widespread cell death, severe fibrosis, and susceptibility to liver damage, including a failure to respond to PH or chemically mediated hepatocyte injury. These phenotypes are associated with a failure to induce genes which are occupied by H3K27me3 in wild-type (WT) livers (10, 11). Interestingly, PRC2 was also implicated in the acquisition of hepatocyte identity, suggested by the finding that *Ezh1 ^−/−^ /Ezh2^HepKO^*postnatal mice had premature activation of hepatocyte identity genes (12), which was further supported by other studies showing that Ezh2 deficiency early in development reduces biliary cell generation (10, 14).

In addition to the role of PRC2 targets in cell identity, the interesting finding that loss of H3K27me3 in hepatocytes causes liver damage, fibrosis and a loss of sex-specific hepatic gene(11–13) expression is not fully understood. Some canonical fibrogenic genes, including *Tgfb1*, *Fbn1*, *Fstl1* and collagen, loose bivalency in *Ezh1 ^−/−^/Ezh2^HepKO^*livers, are shifted to active chromatin and upregulated (12, 13, 16), suggesting this could promote fibrosis in this model. An alternative hypothesis is that H3K27me3 loss causes hepatocyte dysfunction leading to death, which then triggers the well-established cascade of immune infiltration and fibrosis(17). It remains unclear how deregulation of PRC2 leads to liver damage is not clear. A caveat to these studies is that they rely on whole-body deletion of Ezh1 combined with hepatocyte-specific loss of Ezh2, making it difficult to distinguish hepatocyte-intrinsic effects from contributions of other cell types. Importantly, while Ezh1 can compensate for Ezh2 loss in most settings, this does not apply to all PRC2 targets (18, 19). Therefore, the phenotype of *Ezh1 ^−/−^/Ezh2^HepKO^* mice could be due, in part to Ezh1 deficiency in non-hepatocyte populations.

To address the hepatocyte intrinsic mechanisms by which PRC2 alters liver homeostasis and regeneration, we targeted Embryonic Ectoderm Development (Eed) in post-natal hepatocytes (*Eed^HepKO^*). Eed functions as a scaffolding protein, stabilizing PRC2 (18, 20). Full-body knockout of *Eed* is lethal in zebrafish (21) and mice (22), and tissue-specific Eed knockout has emerged as a useful tool to study the effect of Eed in several contexts (18, 23–26). Importantly, EED has emerged as a druggable target for epigenetic therapy, highlighting the importance of understanding how EED loss impacts different cell types to predict the tissue-level consequences of pharmacologic EED inhibition (27–30). We found that H3K27me3 was eliminated in all hepatocytes of young adult *Eed^HepKO^* mice. While these mice appeared phenotypically normal, they had a reduced liver size associated with liver injury, cell death, proliferation, and fibrosis, similar to, but less severe than, Ezh1/2 deficient mice. We found that all genes marked with H3K27me3 in WT livers were induced in *Eed^HepKO^* mice, and these were further upregulated following PH. *Eed^HepKO^* mice had a lower survival following PH, but those that did survive had increased liver mass earlier than WTs. We conclude that examining PRC2 loss specifically in hepatocytes is necessary in order to distinguish hepatocyte-intrinsic regulatory responses from secondary effects arising from other liver cell types.

## Results

### *Eed* knockout depletes H3K27me3 in hepatocytes

We generated mice with a hepatocyte-specific knockout of *Eed* by crossing the Albumin:Cre transgenic line to a floxed allele of *Eed* with *loxP* sequences flanking exons 3 and 6 (31). Offspring were inbred to obtain *Eed^HepKO^*mice which had depleted Eed assessed by quantitative PCR (Figure 1A), Western blot analysis (Figure 1B), and immunofluorescence (Figure 1C) of the liver, as well as complete loss of H3K27me3 (Figure 1D). Moreover, from 2-month-old male *Eed^HepKO^* mice showed no gross phenotypic abnormalities compared to WT counterparts (Figure 1E) and continued to gain weight through 40 weeks of age (Figure 1F).

**Figure 1.**
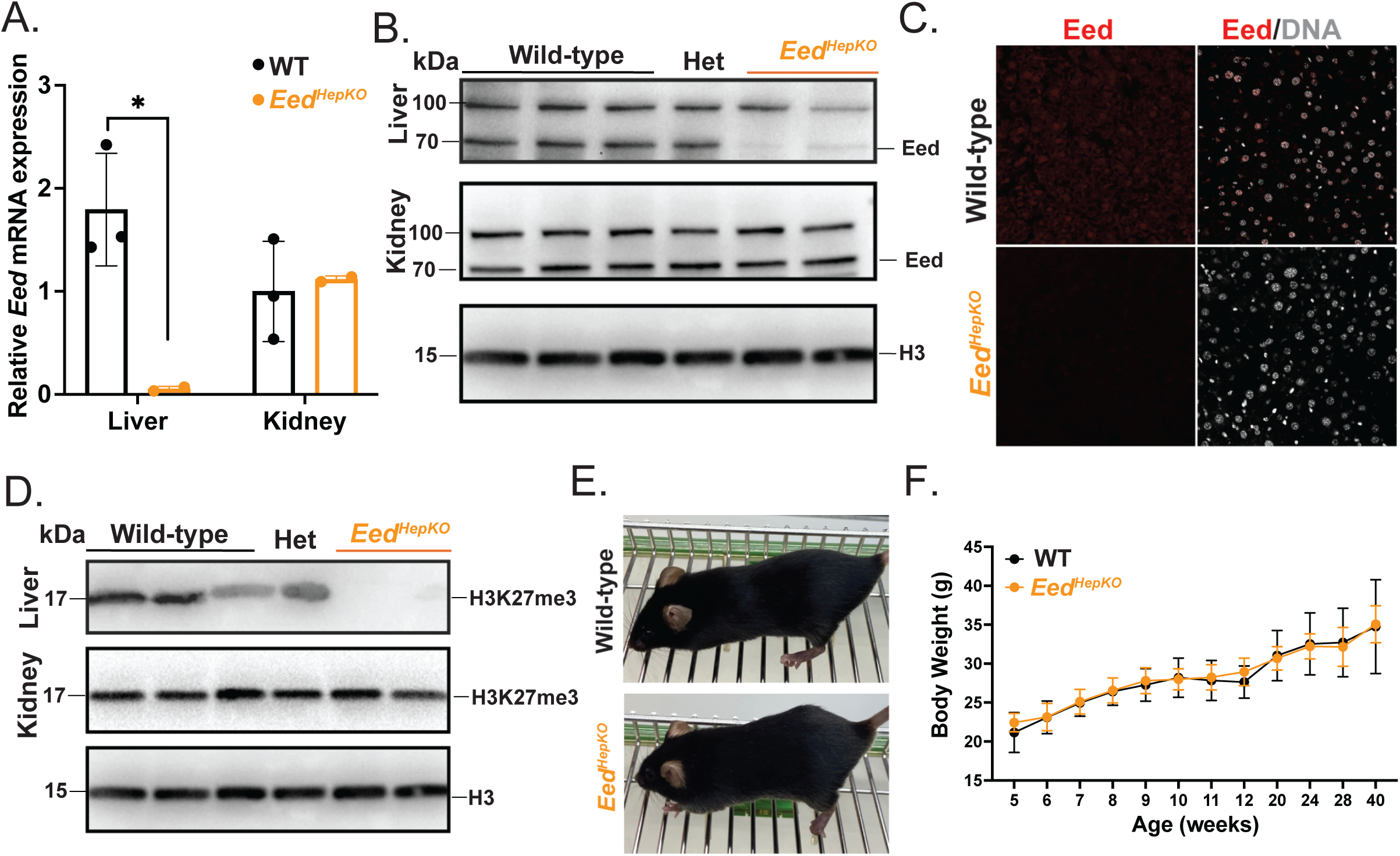
Hepatocyte-specific loss of EED is well-tolerated. **A.** qPCR analysis of *Eed* mRNA expression in 2-month-old, male WT and *Eed^HepKO^* mouse livers and kidneys. **B.** Western Blot analysis comparing the presence of EED in 2-month-old, male WT, *Eed^Het^*, and *Eed^HepKO^* mouse livers. **C.** Immunofluorescent imaging of EED and nuclei in 2-month-old, male WT and *Eed^HepKO^* mice. **D.** Western Blot analysis comparing the presence of H3K27me3 in 2-month-old, male WT, *Eed^Het^*, and *Eed^HepKO^* mice livers and kidneys. **E.** Representative images of 2-month-old, male WT and *Eed^HepKO^* mice. **F.** Body weight measurements of male WT and *Eed^HepKO^* mice between 5 and 40 weeks of age. Data are representative of at least two independent experiments with n = 2-3 (A-D), n = 5-6 (F) per group. Values are presented as mean ± SD and statistical significance was determined using an unpaired two-tailed t-test with Welch’s correction (A) and Two-way ANOVA with Bonferroni’s multiple comparisons test (F). Significance is indicated as *p<0.05, **p<0.01, ***p<0.001, ****p<0.0001.

### *Eed* loss reduces liver size and changes hepatic architecture

We evaluated the effect of Eed loss on the liver, and found 2-month-old male *Eed^HepKO^* mice livers were smaller than WT (Figure 2A-B), though there was no apparent change in gross liver appearance, in comparison to the strong fibrotic phenotype observed in *Ezh1 ^−/−^/Ezh2^HepKO^*(11, 12). Histological analysis of the livers revealed multiple abnormalities (Figure 2C), including increased number of blood vessels (Figure 2D), ductular reactions (20%), biliary fibrosis (20%), proliferation detected by increased number of mitotic figures (20%), and signs of inflammation (60%), none of which pathologies were present in WT controls (Figure 2E). Therefore, loss of PRC2 specifically in hepatocytes causes changes to both hepatocytes and other hepatic cell types.

**Figure 2.**
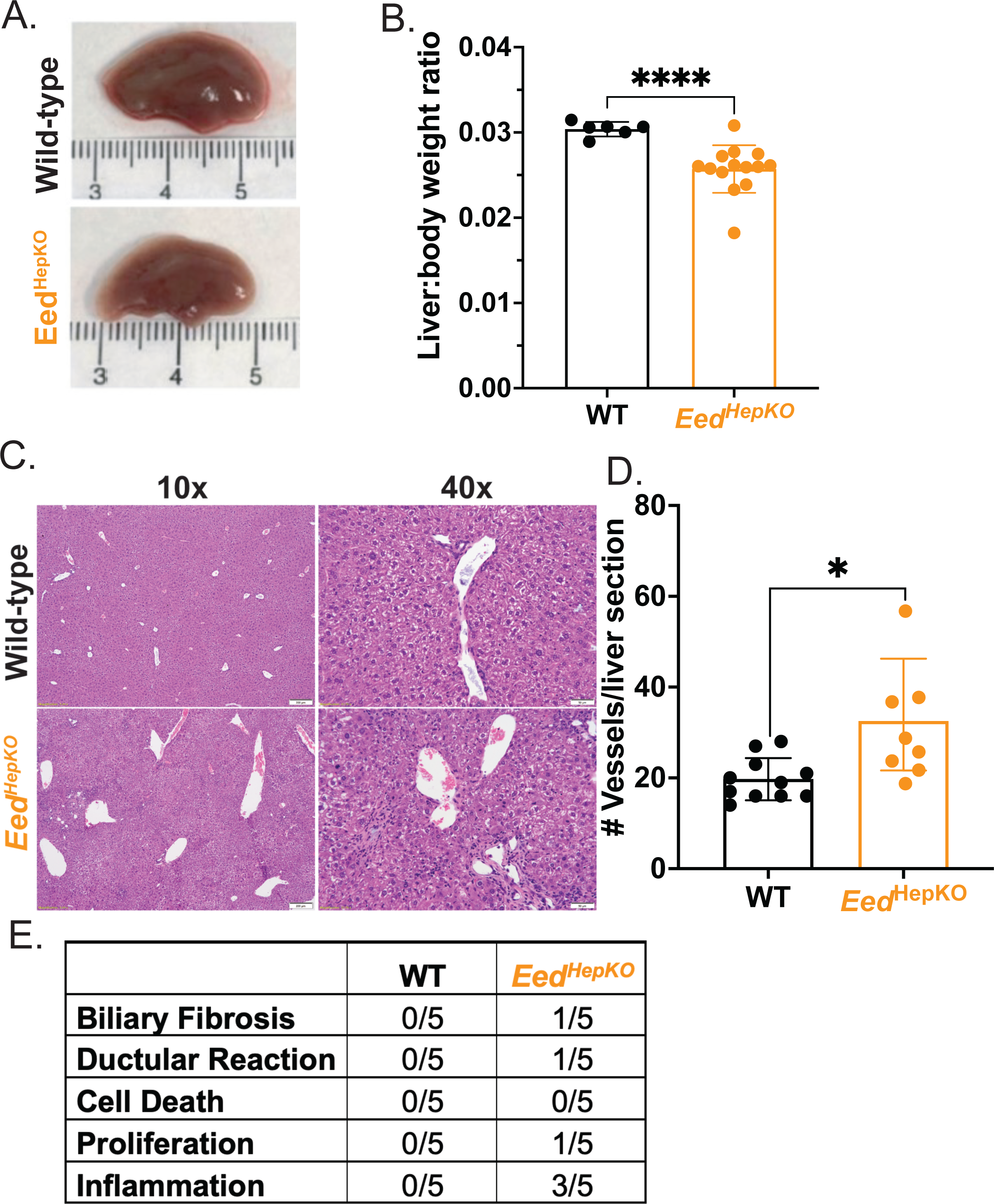
Hepatic loss of *Eed* reduces liver size and increases vascular structures. **A.** Representative images of 2-month-old, male WT and *Eed^HepKO^* mouse livers. **B.** Body-to-liver weight ratios between 2-month-old, male WT and *Eed^HepKO^* mice. **C.** Representative H&E-stained histological images of 2-month-old, male WT and *Eed^HepKO^* mouse livers at 10X and 40X magnification. **D.** Quantification of number of vessels and ducts present in randomly selected 40X magnification images of WT and *Eed^HepKO^* mice. **E.** Table of pathologies present in the WT and *Eed^HepKO^* mice. Data are representative of at least two independent experiments with n = 6-14 (B), n = 8-11 (D) per group. Values are presented as mean ± SD and statistical significance was determined using an unpaired two-tailed t-test with Welch’s correction (B,D). Significance is indicated as *p < 0.05, **p < 0.01, ***p < 0.001, ****p < 0.0001.

### *Eed^HepKO^* changes the hepatic transcriptome

We predicted that the loss of PRC2 activity in hepatocytes had a direct effect on H3K27me3-regulated genes, and, as a response to the phenotype caused by these changes, other genes would become deregulated through an indirect mechanism. Bulk RNA sequencing of WT and *Eed^HepKO^* livers from 2-month-old male mice revealed substantial differences, as shown by principal component analysis (Figure 3A), with 2,566 differentially expressed genes (DEGs) upregulated and 411 downregulated compared to WT livers (log_2_ fold change +/−1; padj <0.05; Figure 3B; Supplemental Table 1). Gene ontology (GO) analysis showed upregulated pathways were involved in tissue development and morphogenesis and downregulated genes were enriched in key hepatocyte metabolic and glycolytic pathways (Figure 3C). Gene set enrichment analysis further identified induction of epithelial to mesenchymal transition, inflammation pathways and apoptosis (Figure S1A).

**Figure 3.**
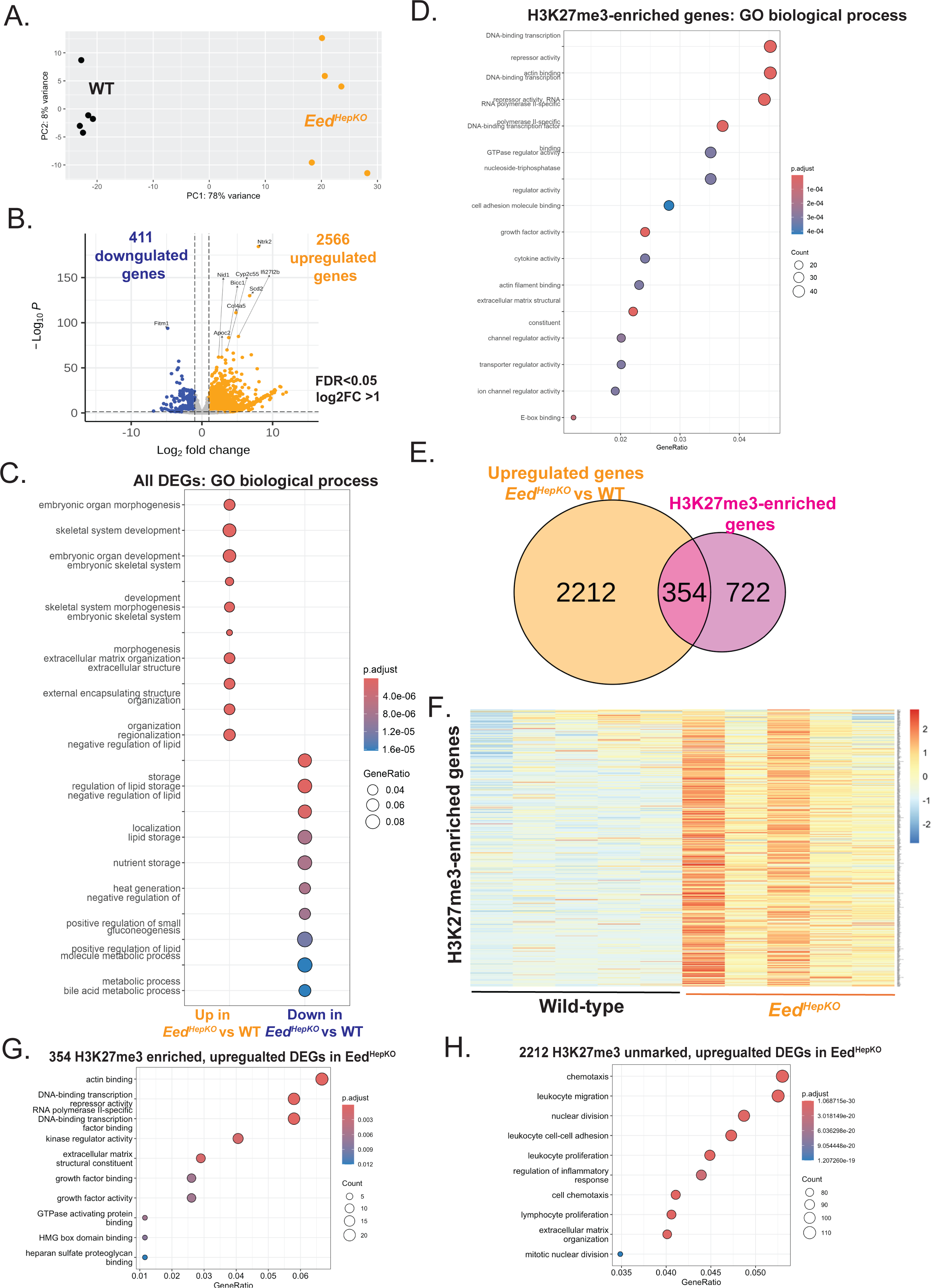
Deregulation of H3K27me3 enriched genes in *Eed^HepKO^* livers. **A.** Principal component analysis (PCA) of liver transcriptomes from 2-month-old male WT (WT) and *Eed^HepKO^* mice. Each point represents an individual biological sample (n = 5 mice per group). **B.** Volcano plot showing differentially expressed genes (DEGs) between WT and *Eed^HepKO^* mice (FDR < 0.05, [log_2_ fold change] > 1). Upregulated genes are shown in orange and downregulated genes in blue. **C.** Gene Ontology (GO) pathway enrichment analysis of genes upregulated and downregulated in *Eed^HepKO^* mice relative to WT controls**. D.** GO pathway enrichment analysis of all known H3K27me3-enriched genes. **E.** Venn diagram showing the overlap between genes upregulated in *Eed^HepKO^* mice and known H3K27me3-enriched genes (354 overlapping genes). **F.** Heatmap displaying expression levels of H3K27me3-enriched genes in livers from WT and *Eed^HepKO^* mice. **G.** GO pathway enrichment analysis of genes that are both upregulated in *Eed^HepKO^* mice and annotated as H3K27me3-enriched. **H.** GO pathway enrichment analysis of upregulated genes in *Eed^HepKO^* mice that are not annotated as H3K27me3-enriched. Data were analyzed using Deseq2, a gene is differentially expressed when log_2_FC (Fold change) > 1 and FDR (False discovery rate) <0.05.

We next investigated which genes were regulated by H3K27me3 by evaluating the 1,076 genes we previously described were occupied by H3K27me3 in promoters of WT livers ((4); i.e. H3K27me3-enriched. GO analysis shows that these genes are involved in a variety of cellular functions including regulation of transcription, growth factor activity, and cell adhesion (Figure 3D; Supplemental Table 2). Of the 2,566 upregulated DEGs in *Eed^HepKO^* livers, 354 were H3K27me3-enriched in WT mice (Figure 3E), but when we assessed expression of all H3K27me3-enriched genes, we see that nearly all were either upregulated or nearly unchanged by the loss of PRC2 activity (Figure 3F), even if not all reached statistical significance. The upregulated DEGs that were H3K27me3-enriched function in pathways associated with transcription and growth factor responses (Figure 3G) and transcription (Figure S1B). We assume these are PRC2 targets that could be the cause of the *Eed^HepKO^* phenotype, and neither the cell cycle nor most of the fibrogenic genes were enriched in this geneset. In contrast, the upregulated genes that were not H3K27me3-enriched functioned in cell cycle, inflammation and the extracellular matrix (Figure 3H). This suggests that these processes were changed in response to the primary defects caused by PRC2 deficiency in hepatocytes.

### *Eed^HepKO^* liver transcriptome resembles that of *Ezh1 ^−/−^/Ezh2^HepKO^*livers

The phenotype of *Eed^HepKO^* mice is reminiscent of the *Ezh1 ^−/−^/Ezh2^HepKO^* mice and we further investigated transcriptomes of these two models. While there were 1,226 DEGs shared between both samples, there were 1,751 and 1,250 DEGs unique to the *Eed^HepKO^*and *Ezh1 ^−/−^/Ezh2^HepKO^* samples, respectively (Figure 4A; Table S3), and the expression of genes in the unified set of genes were found to have the same expression pattern, albeit with subtle differences (Figure 4B). DEGs common to both sets were related to cell differentiation and metabolic processes (Figure 4C). DEGs unique to the *Ezh1 ^−/−^/Ezh2^HepKO^* mice were enriched in functions related to glycolysis and cell respiration (Figure 4D), while DEGs unique to the *Eed^HepKO^* mice were primarily involved in cell cycle regulation, mitosis, DNA replication and other cellular processes. Several studies have shown that fibrogenic genes are upregulated in *Ezh1 ^−/−^/Ezh2^HepKO^*livers and we compared the expression of these genes to *Eed^HepKO^*samples which showed that while all were induced, the magnitude of induction was less than in the double knockout and only a few (*Lox*, *Tgfb1*, *Tgfb2*, *Vim*, *Col4a1*) were likely H3K27me3 targets (Figure 4F). Therefore, while both models have PRC2 loss-of-activity transcriptional responses indicating liver damage and fibrosis, the effects are not identical, and the *Eed^HepKO^* mice have a specific deregulation of genes involved in proliferation, these are not likely to be a direct result of loss of HeK27me3 (Figure 3G). Together, these data demonstrate that the transcriptional phenotype of *Eed* loss in hepatocytes has some shared features *Ezh1 ^−/−^/Ezh2^HepKO^*loss, but with several distinctions and a less pronounced effect on genes involved in fibrosis.

**Figure 4.**
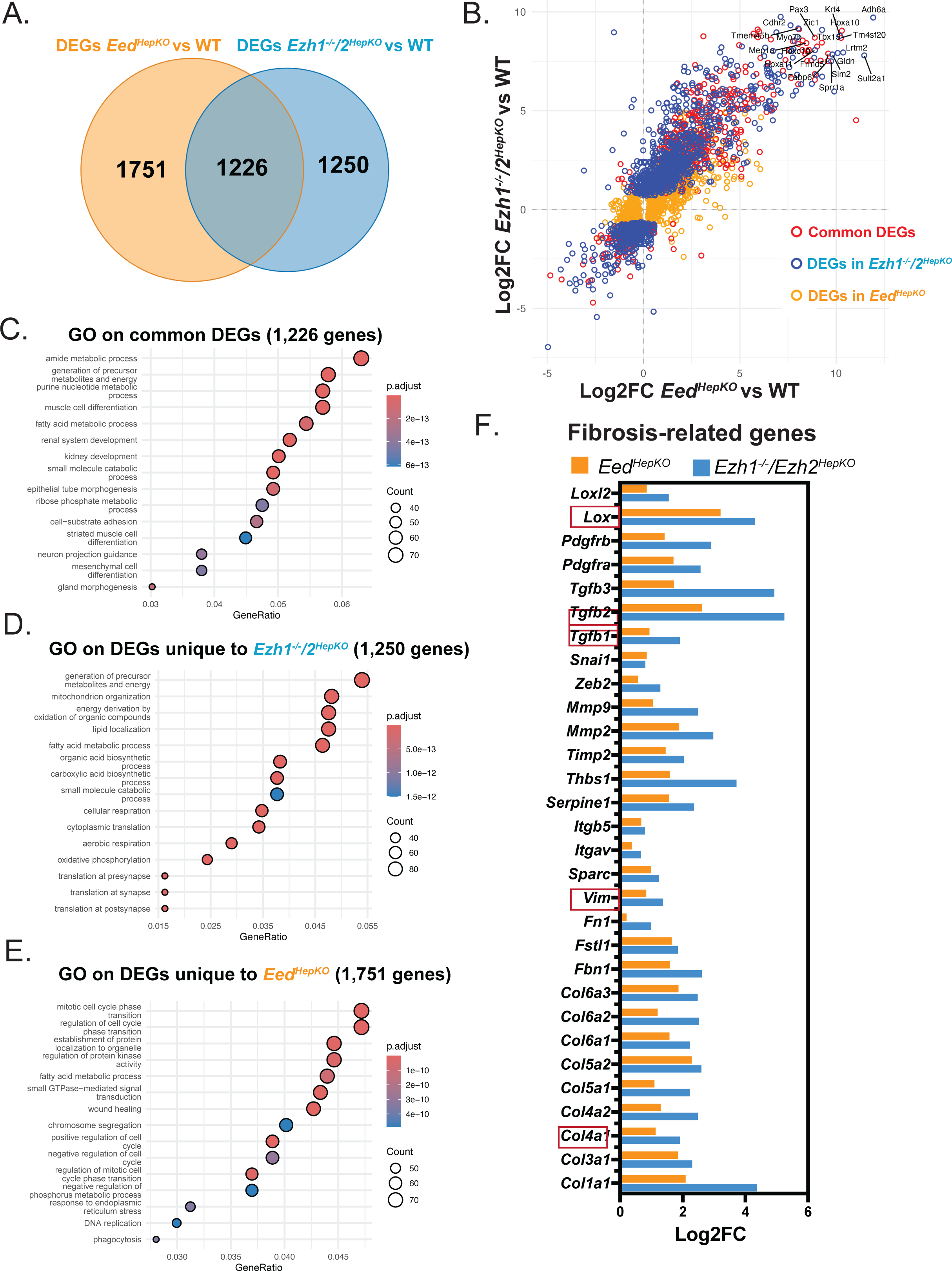
Identification of common and unique genesets deregulated in *Eed^HepKO^* and *Ezh1/2^HepKO^* livers. **A.** Venn diagram showing the overlap of differentially expressed genes (DEGs) between *Eed^HepKO^* and *Ezh1^−/−^/2^HepKO^* mouse livers (1,226 common genes) (*Ezh1^−/−^/2^HepKO^* dataset: *GSE118757*). **B.** Crossplot comparing log_2_FC (fold change) of all DEGs in *Eed^HepKO^* versus WT (WT) and *Ezh1^−/−^/2^HepKO^* versus WT livers. The x-axis represents log_2_FC in *Eed^HepKO^* versus WT and the y-axis represents log_2_FC in *Ezh1^−/−^/2^HepKO^* versus WT. Genes differentially expressed in both models are shown in red, genes uniquely differentially expressed in *Ezh1^−/−^/2^HepKO^* livers in blue, genes uniquely differentially expressed in *Eed^HepKO^* livers in orange. **C-E.** Gene Ontology (GO) pathway enrichment analysis of genes differentially expressed in both *Eed^HepKO^* and *Ezh1^−/−^/2^HepKO^* livers (C), genes uniquely differentially expressed in *Ezh1^−/−^/2^HepKO^* livers (D), and genes uniquely differentially expressed in *Eed^HepKO^* livers (E). **F.** Bar plots showing log_2_FC of fibrosis-associated genes in *Eed^HepKO^* (orange) and *Ezh1^−/−^/2^HepKO^* (blue) mouse livers.

### Hepatic EED loss increases liver damage, cell proliferation, and fibrosis

We investigated phenotypes associated with the transcriptional changes observed in *Eed^HepKO^* mice as nearly all liver damage-associated genes were upregulated (Figure 5A). Serum levels of alanine aminotransferase (ALT) and aspartate aminotransferase (AST) levels were significantly increased roughly three-fold in *Eed^HepKO^* mice, reflecting liver damage (Figure 5B). This was accompanied by an increase in TUNEL-labeled cells as an indicator of cell death in *Eed^HepKO^* livers (Figure 5C). Consistent with the upregulation of genes involved in fibrosis, Sirius Red staining was significantly increased in *Eed^HepKO^* livers (Figure 5D and S2), as were all of the canonical fibrosis-related genes, the majority of which were not marked by H3K27me3 in WT livers (Figures 5E).

**Figure 5.**
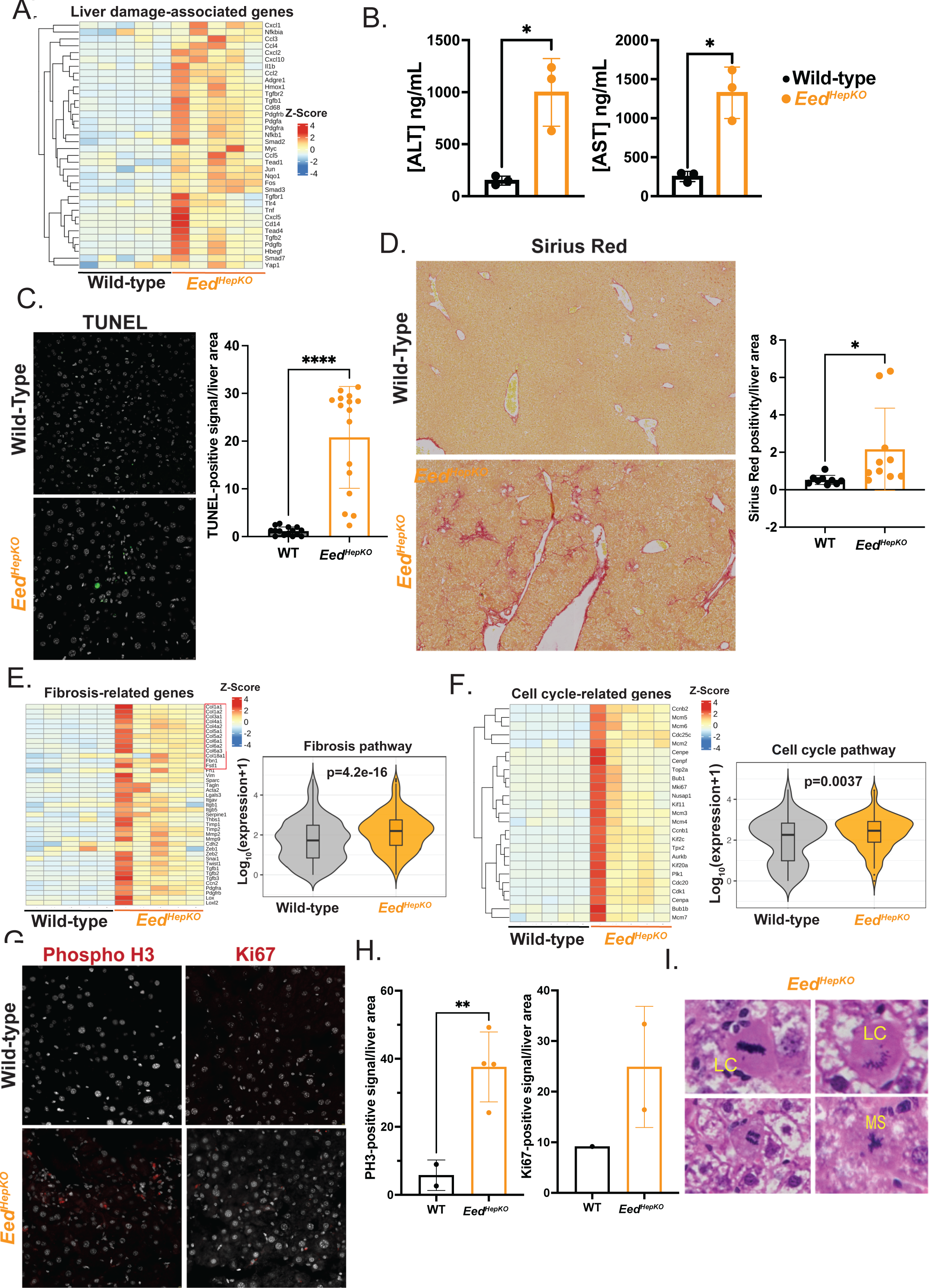
Hepatic EED loss causes liver damage, increased cell proliferation and fibrosis. **A.** Heatmap of liver damage-associated genes in 2-month-old male WT (WT) and *Eed^HepKO^* mice. **B.** Serum alanine aminotransferase (ALT) and aspartate aminotransferase (AST) concentration measured by ELISA in 2-month-old male WT (WT) and *Eed^HepKO^* mice. **C.** Immunofluorescence analysis of hepatocyte apoptosis in liver cryosections from 2-month-old WT and *Eed^HepKO^* mice using TUNEL staining. Representative images and quantification of TUNEL-positive cells per liver section are shown. **D.** Representative Sirius Red/Fast Green staining of liver sections from 2-month-old WT and *Eed^HepKO^* mice (40× magnification) with quantification of collagen-positive area per section. **E**. Heatmap showing z-score normalized expression of fibrosis-associated genes in WT and *Eed^HepKO^* livers, with violin plots displaying log10 expression of fibrosis pathway genes. **F.** Heatmap and violin plot analysis of cell-cycle–associated gene expression in WT and *Eed^HepKO^* livers. **G.** Representative immunofluorescence images and quantification of hepatocyte proliferation markers phospho-histone H3 (pH3) and Ki67 in liver sections from WT and *Eed^HepKO^* mice. **H.** Heatmap showing z-score normalized expression of Liver damage-associated genes in WT and *Eed^HepKO^* livers. Data are representative of at least two independent experiments with n = 3 (A), n = 5 (B), n = 4 (C), n = 5 (D-E, H), and n = 1-4 (F) mice per group. Values are presented as mean ± SD and statistical significance was determined using an unpaired two-tailed t-test with Welch’s correction (A-B,F). Significance is indicated as *p < 0.05, **p < 0.01, ***p < 0.001, ****p < 0.0001. **I.** Representative hematoxylin and eosin (H&E)–stained liver sections showing mitotic figures in *Eed^HepKO^* baseline livers

To further investigate the proliferation status indicated by the finding that cell cycle pathways were induced in *Eed^HepKO^* livers (Figures 3H, 4E) we analyzed the expression of the entire cell cycle hallmark geneset (GO:0000086) and found all to be induced (Figure 5F). This was reflected in the detection of increased markers for mitotic cells (phospho-histone H3) and proliferation, Ki67 (Figures 5G and 5H). Interestingly, histological analysis of the mitotic figures showed several that appeared to be abnormal, with lagging chromosomes and multipolar spindles (Figure 5I). This suggests that PRC2 deficiency in hepatocytes causes upregulation of most genes suppressed by H3K27me3 in WT livers, leading to hepatocyte damage and death which induce proliferation and pro-fibrogenic signals a secondary response.

### *Eed^HepKO^* mice have divergent responses to PH

We hypothesized that loss of H3K27me3 on pro-regenerative genes allows them to be activated following a regenerative stimuli, which is consistent with the finding that H3K27me3 levels were reduced during regeneration of WT mice (4) (Figure 6A) and repatterning of H3K27me3 from promoters is associated with enhanced liver regeneration (9). However, the finding of liver damage and fibrosis in *Eed^HepKO^*livers suggests that, like *Ezh1 ^−/−^/Ezh2^HepKO^* mice (11, 13), they may be unable to mount a regenerative response, even if the proregenerative genes were able to be activated. To test this, we performed PH on young *Eed^HepKO^* and WT male mice and assessed survival, restoration of liver mass and the transcriptional response. In comparison to 100% survival of WT, only 64% of the *Eed^HepKO^* mice survived PH (Figure 6B). However, those *Eed^HepKO^* mice that did survive, recovered liver mass faster than WT controls, as measured by their liver/body weight ratio (Figure 6C).

**Figure 6.**
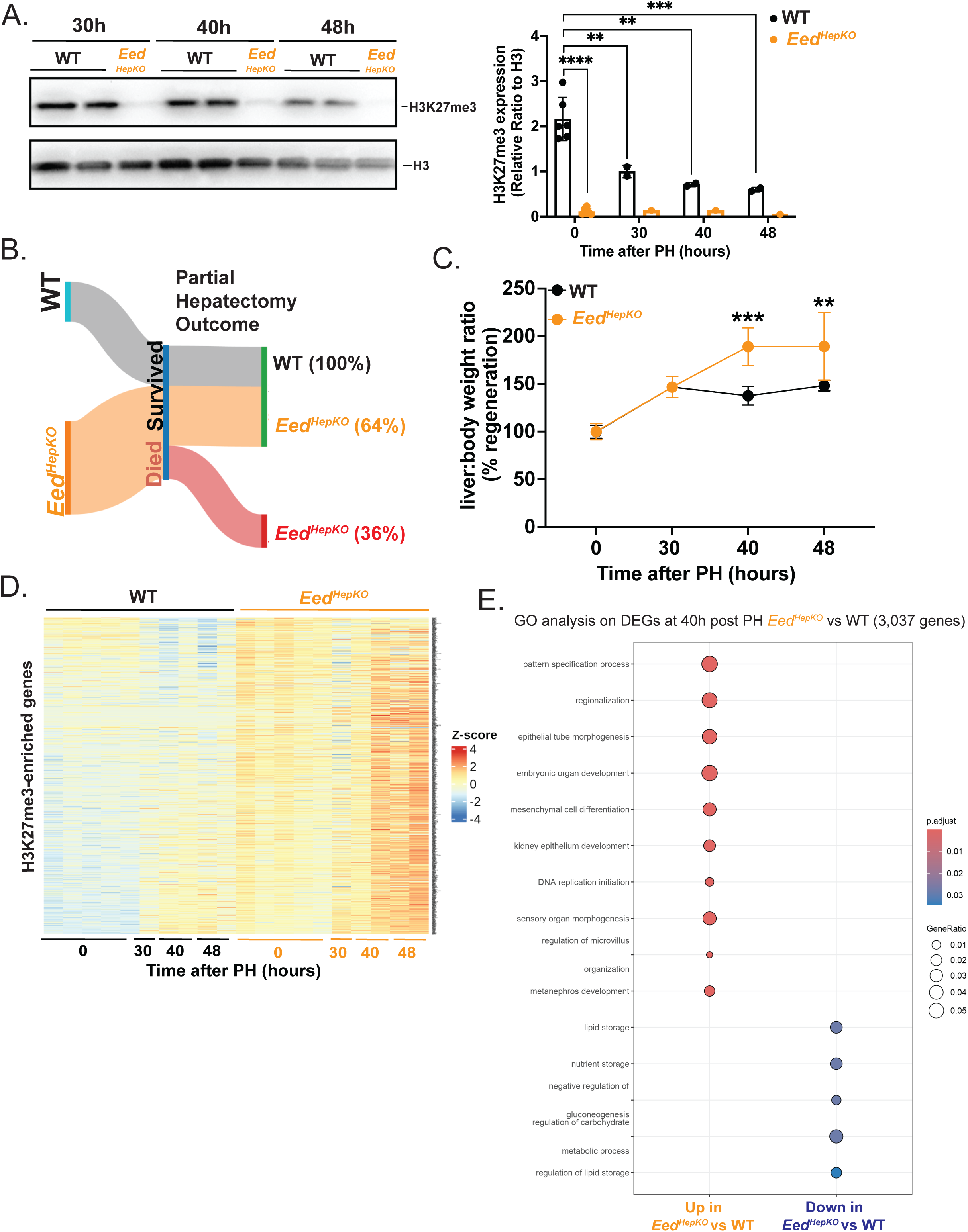
*Eed^HepKO^* mice show a binary response to PH. **A.** Western blot analysis of H3K27me3 and total histone H3 in liver extracts from WT (WT) and *Eed^HepKO^* mice at baseline and 30, 40, and 48 hours after two-thirds partial hepatectomy (PH), with quantification of H3K27me3 levels normalized to histone H3. **B.** Visualization of PH outcome (survival and death) of WT and *Eed^HepKO^* mice following two-thirds PH. **C.** Quantification of liver regeneration following PH, measured as liver-to-body weight ratio at indicated time points. **D.** Heatmap showing z-score normalized expression of H3K27me3-enriched genes in WT and *Eed^HepKO^* livers before and after PH (30 h, 40 h, and 48 h) determined by bulk RNA sequencing. **E.** Gene Ontology (GO) pathway enrichment analysis of differentially expressed genes (DEGs) at 40 hours post-PH in *Eed^HepKO^* mice relative to WT controls 40 hours post-PH. Data are representative of at least two independent experiments with n = 1-6 (A), n = 10-14 (B), n = 3-4 (C), and n = 5 (D-E) mice per group. Values are presented as mean ± SD, and statistical significance was determined using an unpaired two-tailed t-test with Welch’s correction (A) and Two-way ANOVA with Bonferroni’s multiple comparisons test (C). Significance levels are indicated as *p < 0.05, **p < 0.01, ***p < 0.001, ****p < 0.0001. Transcriptomics data were analyzed using Deseq2, a gene is differentially expressed when log_2_FC (Fold change) > 1 and FDR (False discovery rate) <0.05.

Bulk RNAseq analysis of WT and *Eed^HepKO^* mice after PH showed that while some H3K27me3-enriched genes were downregulated during regeneration in WT livers, most were induced. The profile in *Eed^HepKO^*livers was strikingly different, with all these targets being overexpressed in uninjured livers, then downregulated at 30 hours after PH, and nearly all were induced at 40 and 48 hours after PH, the time points when hepatocytes proceed from S phase to mitosis (Figure 5D). Interestingly, at 40 hours, the time point when many pro-regenerative genes are upregulated in controls (4, 9) the DEGs between *Eed^HepKO^* and WT were mostly involved in development and cell division, whereas the downregulated genes were canonical hepatocyte metabolic functions (Figure 5E). This suggests that PRC2 loss causes hepatocyte damage which impairs liver regeneration in some mice, while those that survive are able to compensate, with upregulated proliferative genes and increased liver mass.

## Discussion

Liver development and regeneration rely on tightly regulated gene expression patterns generating cell identity and function. Studies from many systems point to an important role of PRC2 mediated regulation of genes that promote regeneration (5–8, 32, 33). Given the unique regenerative capacity of the young liver, several studies have investigated whether PRC2 mediated H3K27me3 gene regulation is important for liver regeneration. These approaches using a whole-body knockout of Ezh1 and hepatocyte specific Ezh2 due to the effective but not complete redundancy of these proteins (15, 18, 19, 24) and therefore it is possible that some of the phenotype observed are attributed to Ezh1 loss in other cell types. We addressed this by generating mice with hepatocyte specific knockout of Eed which provides a simplified model to understand PRC2 function in the liver. We found that, like *Ezh1 ^−/−^/Ezh2^HepKO^* mice, *Eed^HepKO^* caused liver damage, cell death, fibrosis and compromised response to regenerative stimuli. However, the phenotype was less pronounced than that described for *Ezh1 ^−/−^/Ezh2^HepKO^* model, which develop severe fibrosis. This is accompanied by induction of nearly all genes proposed to be regulated by H3k27me3 based on promoter occupancy as well as the deregulation of over 2,000 genes which are not H3K27me3 enriched. This suggests that the deregulation of H3K27me3 genes is the primary defect causing hepatocyte damage and death, which then results in the deregulation of other processes, such as cell proliferation, inflammation and fibrosis as a response. It is not clear which processes deregulated in this model are responsible for cell damage and dysfunction.

The most striking phenotype in *Eed^HepKO^* is the liver damage in the absence of injury, as indicated by increases in serum ALT and AST levels, as well as the presence of TUNEL-positive cells and the upregulation of damage-associated genes. Fibrosis was also evident, and was accompanied by significant upregulation of fibrosis-associated genes. In contrast to the normal adult liver, which contains very few actively dividing cells under homeostatic conditions, there were numerous proliferating cells along with increased expression of cell division markers and cell cycle genes in *Eed^HepKO^* livers. Interestingly, many of the mitotic figures had evidence of mitotic defects, including lagging chromosomes and multipolar spindles. Early work on PRC deficiency in *Drosophila* reported multiple mitotic defects (34) and a more recent study found that Ezh2 loss induced a similar phenotype in cancer cells (35). One possible mechanism is that mitotic defects caused by PRC2 loss lead to hepatocyte dysfunction and death, triggering compensatory regeneration. The resulting hepatocyte injury may then promote inflammation and fibrosis as secondary responses.

We found that Eed loss caused a small for sized liver in adults, similar to other studies with PRC2 deficiency in the liver (10–12)and this may be a result of hepatocyte loss or to a developmental defect. Other studies have shown that PRC2 loss in the liver has a profound effect on development. Deletion in postnatal livers results the activation of genes which are typically activated in adults (12), which was interpreted as a premature maturation of hepatocytes in this model. Other studies show that loss of Ezh2 earlier in development has a different effect, causing a reduction in hepatic progenitor cells and a failure of those progenitors that do form to differentiate into hepatic and biliary lineages (10). There is strong evidence that genes dictating the hepatocyte and biliary lineages are bivalent in early progenitors and then transition to becoming active in biliary cells and retaining their plasticity, while the loss of the bivalent mark H3K4me3 restricts the plasticity of hepatocytes, and Ezh2 knockout increases the number of biliary cells (14). Further investigation into the cell composition of the *Eed^HepKO^* livers will enable determination of whether biliary reaction and vascular defects represent a patterning defect during liver development or are a reaction to the hepatocyte dysfunction and death.

PRC2 activity is required for embryo viability (16, 21) but several studies have found that tissue specific loss reveals distinct roles in a cell type specific fashion. For instance, in hematopoietic stem cells, Eed loss causes depletion of these cells in adults, but fetal maturation appears normal (18). Eed knockout in Schwann cells impairs proliferation and induces the cell cycle inhibitors (36) while Eed mediated depletion of PRC2 in hair follicle stem cells has no effect on regeneration. This points to the importance of using cell specific models to dissect the role of PRC2 in each cell type. Our finding of a similar yet less severe phenotype in the *Eed^HepKO^*mice compared to *Ezh1 ^−/−^/Ezh2^HepKO^* livers, including increased number of blood vessels, biliary fibrosis, actively dividing hepatocytes, inflammation, and fibrosis in uninjured livers. Moreover, the similar gene expression changes observed between *Eed^HepKO^* to *Ezh1 ^−/−^/Ezh2^HepKO^*, even though DEGs only partially overlapped, both strains presented significantly upregulated genes relating to fibrosis. One explanation is that Ezh1 deficiency in hepatic stellate cells or immune cells may not be fully compensated by Ezh2, allowing PRC2 loss in these non-hepatocyte populations to contribute to the phenotype. However, because our analysis compares *Eed^HepKO^* mice with phenotypes reported in prior studies, differences in non-genetic factors such as microbiome composition, genetic background, or housing conditions could also contribute to the variation between models.

Regardless of the subtle differences between different models of hepatic PRC2 loss, we hypothesize that the baseline level of liver injury limits the regenerative capacity. Consistent with this, we report that 1/3 of *Eed^HepKO^*mice died between 2-30 hours after PH, suggesting they were unable to mount a regenerative response. In patients, several factors contribute to liver failure following PH, including ischemia, portal hypertension, failure to mount a proliferative response, hepatocyte cell death and steatosis (37). Although we did not find evidence of steatosis however, there was noted inflammation, especially periportally and pericentrally, raising the possibility that vascular defects which may restrict blood flow in the regenerating liver, leading to ischemia or failed proliferation.

Interestingly, the *Eed^HepKO^* mice that survived PH showed increase in liver mass at time points earlier than WT mice. While this was accompanied by an upregulation of PRC2-associated genes, the same genes were also induced in uninjured livers, despite *Eed^HepKO^* mice having small for size livers. Our previous work showed that repatterning of DNA methylation in hepatocytes leads to a redistribution of H3K27me3 from promoters, but this was insufficient to induce gene expression changes in the absence of injury. However, following PH, the genes that lost H3K27me3 were prematurely activated (9). Therefore, while it is possible *Eed^HepKO^* mice are primed for regeneration as many regenerative genes are already open to transcription, other factors, such as edema, could also contribute to the increased liver mass.

These findings confirm and extend previous studies on the role of H3K27me3 and PRC2 in the liver. There is significant interest in epigenetic therapy for a number of diseases (2). For instance, the decline in regenerative capacity during aging (38, 39) is associated with an increase in H3K27me3 levels (8, 40), suggesting the interesting possibility that suppression of pro-regenerative genes by PRC2 can contribute to age related regeneration incompetence, suggesting the exciting possibility that transient alleviation of H3K27me3 mediated repression of proregenerative genes could promote regeneration. However, this should be measured against potential cellular defects caused by such manipulations, as shown by the finding of liver injury and fibrosis in PRC2 deficient mice. Therefore, any approaches to manipulate the epigenome in regenerative medicine requires the precise control of PRC2 function to enhance regenerative capacity.

## Data and Code availability

Raw data (fastq files) and processed files (raw counts) for bulk RNAseq are publicly available on GEO (GSE329873). This paper does not report original code.

## Acknowledgments

This work was supported by the NYUAD Faculty Research Fund (AD188), NIH (2R01DK080789-12A1) to KCS, and Tamkeen under the NYU Abu Dhabi Research Institute Award to the NYUAD Center for Genomics and Systems Biology (ADHPG-CGSB13). All imaging was carried out in the NYUAD Core Technology Platform Imaging facility with expert support by Rashid Razgui. RNAseq bulk libraries were performed by NYUAD Core Technology Platform Imaging with the expert help of Marc Arnoux and bioinformatics assistance from the NYUAD Bioinformatics Core. We are grateful for Sirius Red staining by Elena Magnani, TUNEL staining by Balnur Ibrash and the technical support during collaboration with the Halder Lab (KU-Leuven). We appreciate the expertise of Nizar Drou for assistance with bulk RNAseq. We are thankful to all Sadler lab members for valuable discussions.

## Author contributions

JG, YA, and KCS conceptualized the project, JG, YA, SC, CC, MM, and KCS developed the methodology, JG, YA, CC, MM, KCS carried out the analysis, JG, YA, SC, and MM carried out the investigation, JG, YA, SC, CC, and KCS created the visualization, KCS provided resources, KCS provided supervision, project administration, JG, YA, and KCS wrote the manuscript. JG, YA, CC, and KCS reviewed and edited the manuscript.

## Supplemental information titles and Legend

**Figure S1.** Gene Set Enrichment Analysis of *Eed^HepKO^*mice

**Figure S2.** Sirius red staining of WT and *Eed^HepKO^* mice livers at 3X magnification.

**Supplemental Table S1.** Gene counts from bulk RNAseq analysis of the liver.

**Supplemental Table S2.** Upregulated genes in *Eed^HepKO^*mice.

**Supplemental Table S3.** Aggregated list of upregulated genes in *Ezh1 ^−/−^/Ezh2^HepKO^* mice from Grindheim, et al.

**Supplemental Table S4.** Gene counts from bulk RNAseq analysis of the liver before and after PH in WT and *Eed^HepKO^* mice.

## Materials and Methods

### Generation of *Eed^HepKO^* mice

Mouse husbandry and care was conducted according to the New York University Abu Dhabi (NYUAD) Institutional Animal Care and Use Committee (IACUC) protocol 23-0011A. Mice were weaned at 21-days-old and fed *ad libitum*.

C57BL/6J mice were purchased from The Jackson Laboratory: WT mice, transgenic mice possessing Cre under the expression of the albumen promoter (“Alb:Cre”), and targeted mutant mice with *loxP* sites flanking exons 3 and 6 of the embryonic ectoderm development (*Eed*) gene (“Eed^flox^”; strain #022727). Alb:Cre and Eed^flox^ mice strains were crossed and further inbred to produce “*Eed^HepKO^*” mice that lack functional EED in hepatocytes. Genotyping was performed by Transnetyx, Inc. (Cordova, TN) via polymerase chain reaction on mouse tail clippings obtained when weaned at 21-days-old.

### Quantitative reverse-transcription PCR (qPCR)

RNA was extracted from mouse tissues stored at −80°C using TRIzol (Invitrogen, 15596026) in a Dounce homogenizer. RNA was isolated via column extraction on a QIAGEN RNeasy Mini Kit in tandem with on-column DNase I digestion per the manufacturer’s instructions. RNA was quantified on a Qubit^TM^ fluorometer. RNA samples were reverse-transcribed into cDNA using Quantabio qScript cDNA SuperMix (Quantabio, 95048–025). QPCR was performed using SYBR green (ThermoFisher, 4309155) on a QuantStudio 5 (Applied Biosystems). Samples ran in triplicates and relative expression was quantified by calculating ΔΔCt values against rplp0, the endogenous control, and plotted for visualization in GraphPad Prism 10.

### Partial Hepatectomy

Partial hepatectomy (PH) was performed as described by Mitchell et al. (Mitchell and Willenbring, 2008). Male mice aged 8–12 weeks were anesthetized with isoflurane (5% for induction followed by 1.5–2% for maintenance) between 08:00 am and 12:00 pm to minimize circadian variability. After induction of anesthesia, mice were weighed and placed on a sterile surgical field. A midline abdominal incision (∼3 cm) was made to expose the liver, followed by opening of the peritoneal cavity. The falciform ligament was transected and a two-thirds hepatectomy was performed by ligating and resecting the left lateral and median lobes using 4-0 silk sutures (Assut Sutures, L4553WF). The peritoneal cavity was closed with 4-0 sutures (Assut Sutures, L4553WF), and the skin was closed with wound clips (BrainTree Scientific, EZC APL). Mice were euthanized at 30, 40, or 48 hours following surgery. For measuring the weight of the regenerating liver following surgery, only the intact, non-resected lobes were measured. Tissue samples were snap-frozen in liquid nitrogen and stored at −80 °C for downstream analyses.

### Western blotting

Protein was extracted from mouse tissue stored at −80°C using T-PER^TM^ Tissue Protein Extraction Reagent (Thermo Scientific) supplemented with protease inhibitor cocktail (Roche), and the tissue lysate was cleared via centrifugation. Protein concentration was quantified via Qubit^TM^ assay. Extracted protein was mixed with 4X Laemmli buffer (BioRad), briefly boiled, spun to remove insoluble content, cooled, and loaded into a 10% acrylamide/bisacrylamide denaturing gel. Voltage was initially set to 40V to allow for uniform entry into the gel, at which point the voltage was raised to 100V. Proteins were transferred to a PVDF membrane at 300mA for 1.5 hours on ice. Membrane was blocked for 1 hour at room temperature with TBS (BioRad) supplemented with 0.1% Tween-20 (Sigma Aldrich) containing 5% skim milk (Sigma Aldrich), incubated overnight at 4 °C with 1: 1000 primary antibody diluted in blocking solution (EED, Cell-Signaling, Cat. No. 51673S; H3K27me3, Active Motif, Cat. No. 61017; H3, Santa Cruz, Cat. No. 10809) followed by secondary antibody diluted 1: 2500 in blocking solution (HRP-conjugated anti-Rabbit or anti-Mouse, Promega). Chemiluminescence was revealed with ECL Clarity (BioRad) and imaged with ChemiDoc MP Imaging System (BioRad).

### Immunofluorescence

Mouse liver tissue was resected and fixed in 10% neutral-buffered formalin (Sigma Aldrich) at 4°C for 24-48 hours. Tissue was cryopreserved by incubation in 30% sucrose in PBS overnight, followed by mounting in OCT (Leica Biosystems) and frozen with dry ice. 14 μm cryosections were used for immunofluorescence. Sections were rehydrated with PBS, permeabilized with 1% Tween-20 (Sigma Aldrich) in PBS for 5 minutes, and blocked with 2.5% BSA (Sigma Aldrich) and 0.1% Triton X-100 (Sigma Aldrich) in PBS for 1 hour. Primary antibody was dissolved 1:200 in blocking buffer and incubated for 3 hours. After washing the sections, secondary antibody was dissolved 1:400 in blocking buffer and incubated for 1 hour. Sections were then washed with PBS and mounted in Vectashield containing DAPI (Vector) for imaging. Imaging was performed on a Leica Stellaris 8 Confocal Microscope and quantified with LasX software.

### Histology

Mouse liver tissue was resected and fixed in 10% neutral-buffered formalin (Sigma Aldrich) at 4°C before mounting into paraffin blocks. Sections of the blocks were deparaffinized in xylene, rehydrated, and stained/counterstained with either hematoxylin/eosin or Sirius Red/Fast Green. Stained sections were then dehydrated and mounted. Imaging was performed on a DSX2000 Digital Microscope (Evident Scientific) and quantified with PRECi software. Histopathological analysis was performed by a board certified gastroenterology pathologist

### Alanine aminotransferase/AST serum analysis

Blood was collected from mice via retro-orbital bleeding. After allowing the blood to clot for 30 minutes at room temperature, blood was centrifuged at 1,000 x G for 10 minutes, and the top serum layer was collected and frozen at −80°C for later analysis. ALT and AST concentrations were measured using the Mouse ALT ELISA Kit (Abcam) or the Mouse AST ELISA Kit (Abcam), respectively, and were plotted for visualization with Graphpad Prism 10.

### TUNEL staining

14 μm cryosections of mouse liver tissue were prepared as described above. Cell death was assessed using the In Situ Cell Death Detection Kit, Fluorescein or TMR Red (REF 11684795910/12156792910; Roche). In brief, cryosections were washed with PBS for 30 minutes and permeabilized with 0.1% Triton X-100 (Sigma Aldrich) and 0.1% sodium citrate (Sigma Aldrich) for 2 minutes. Per the manufacturer’s instructions, sections were washed with PBS an additional 2 times, incubated with the kit-provided TUNEL reaction mixture for 60 minutes at 37°C, and washed 3 times with PBS. Sections were mounted in Vectashield containing DAPI (Vector) for imaging. Imaging was performed on a Leica Stellaris 8 Confocal Microscope and quantified with LasX software.

### Sequencing library preparation and bulk RNA-seq analysis

Livers were resected from 2-month-old WT and Eed^HepKO^ mice either before or after partial hepatectomy and stored at −80°C. Tissue was homogenized in 500 ml of TRIzol (Thermo Fisher Scientific) in a Dounce homogenizer. RNA was isolated using column extraction on a QIAGEN RNeasy Mini Kit (Qiagen, 74104) in tandem with on-column DNase I digestion (Qiagen, 79254) as per the manufacturer’s instructions. RNA was quantified on a QubitTM fluorometer and quality-checked on an Agilent TapeStation. 1000 ng of total RNA was subsequently used to prepare RNA-seq library by using TruSeq RNA sample prep kit (Illumina) according to the manufacturer’s instructions. Paired-end RNA-sequencing was performed on a NovaSeq 6000 (Illumina) (CTP-NYUAD, Abu Dhabi, UAE). Sequenced reads were aligned to the mouse genome (mm10), and uniquely mapped reads were used to calculate gene expression. Raw FASTQ sequenced reads where first assessed for quality using FastQCv0.11.5. The reads where then passed through Fastpversion 0.20.0 for quality trimming and adapter sequence removal using the default parameters. Following the quality trimming, the reads were assessed again using FastQC. Data analysis was performed using R program (DESeq2 package). Differentially expressed genes are considered significant when the false discovery rate (FDR or adjusted p-value) < 0.05 and the log_2_ fold change (log2FC) > 1. All RNA-seq analyses were performed using ≥2 biological replicates.

### Statistical analysis

All statistical tests, error bars, and N numbers are reported in the corresponding figure legends with p-values indicated in the figure when significant. All statistical analyses were conducted using GraphPad Prism (GraphPad Software). Image quantification of the immunofluorescence staining was done in FIJI with the Color Thresholding function, where the Threshold Color Space was set to the RGB channel to generate a mask before applying the Threshold Adjuster to select stain-positive pixels.

