## Supplemental Figures for "Hepatocyte Embryonic Ectoderm Development (Eed) Deficiency Causes Liver Injury, Fibrosis, and Impacts Liver Regeneration"

### Supplemental Materials

#### Supplemental Figure S1. Multiple pathways disrupted buy *Eed* loss in hepatocytes.

A. Gene Set Enrichment Analysis of all DEGs in 2 month male ***Eed*$^{HepKO}$ livers**

B. Molecular function by GO analysis of all upregulated DEGs from *Eed*$^{HepKO}$ livers and those marked with H3K9me3 in uninjured WT livers.

**Supplemental Figure S2. *Eed*$^{HepKO}$ induces fibrosis**

Sirius Red staining of WT and *Eed*$^{HepKO}$ liver sections

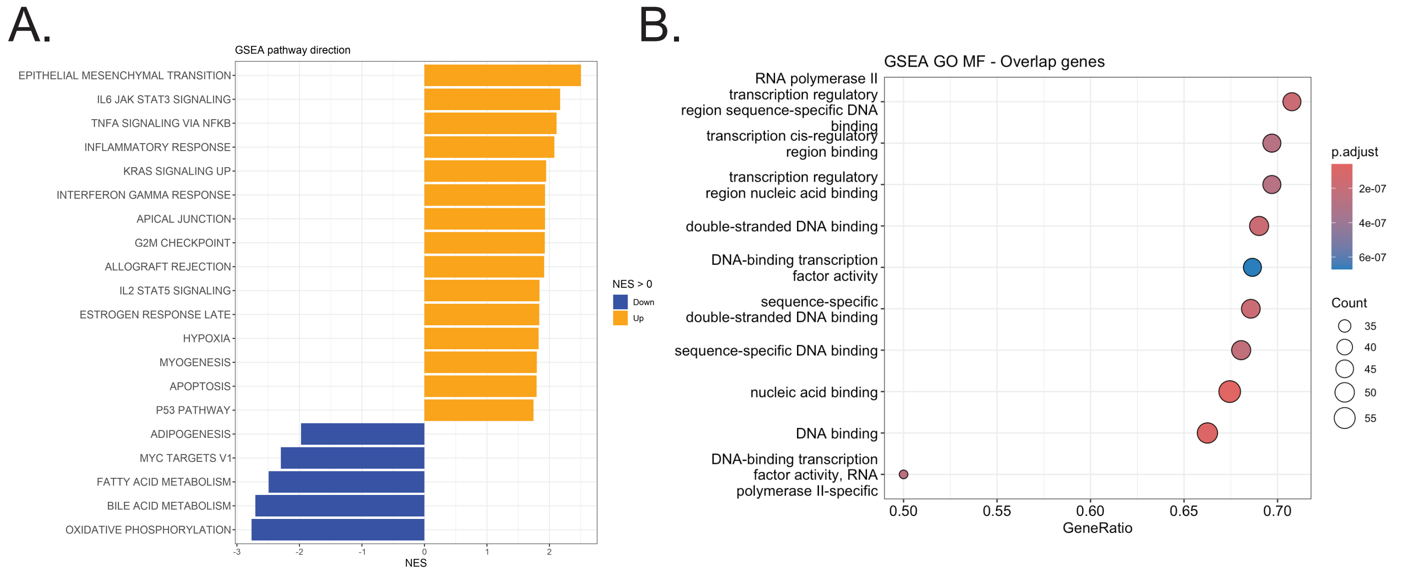
A.
GSEA pathway direction
EPITHELIAL MESENCHYMAL TRANSITION
IL6 JAK STAT3 SIGNALING
TNFA SIGNALING VIA NFKB
INFLAMMATORY RESPONSE
KRAS SIGNALING UP
INTERFERON GAMMA RESPONSE
APICAL JUNCTION
G2M CHECKPOINT
ALLOGRAFT REJECTION
IL2 STAT5 SIGNALING
ESTROGEN RESPONSE LATE
HYPOXIA
MYOGENESIS
APOPTOSIS
P53 PATHWAY
ADIPOGENESIS
MYC TARGETS V1
FATTY ACID METABOLISM
BILE ACID METABOLISM
OXIDATIVE PHOSPHORYLATION
-3
-2
-1
0
1
2
NES
NES > 0
Down
Up
B.
GSEA GO MF - Overlap genes
RNA polymerase II transcription regulatory region sequence-specific DNA binding
transcription cis-regulatory region binding
transcription regulatory region nucleic acid binding
double-stranded DNA binding
DNA-binding transcription factor activity
sequence-specific double-stranded DNA binding
sequence-specific DNA binding
nucleic acid binding
DNA binding
DNA-binding transcription factor activity, RNA polymerase II-specific
0.50
0.55
0.60
0.65
0.70
GeneRatio
p.adjust
2e-07
4e-07
6e-07
Count
35
40
45
50
55

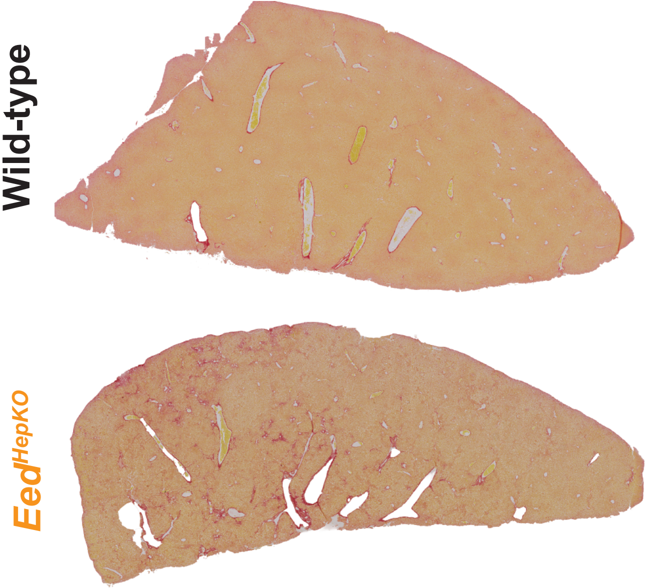
Wild-type
Eed HepKO
